# Developmentally-orchestrated mitochondrial processes prime the selection against harmful mtDNA mutations

**DOI:** 10.1101/646638

**Authors:** Zhe Chen, Zong-Heng Wang, Guofeng Zhang, Christopher K. E. Bleck, Dillon J. Chung, Grey Madison, Eric Lindberg, Christian Combs, Robert S. Balaban, Hong Xu

## Abstract

Although mitochondrial DNA (mtDNA) is prone to mutation and not all conventional DNA repair systems operate in mitochondria, deleterious mutations are exceedingly rare. How the transmission of detrimental mtDNA mutations is restricted through the maternal lineage is debated. Here, we use *Drosophila* to dissect the mechanisms of mtDNA selective inheritance and understand their molecular underpinnings. Our observations support a purifying selection at the organelle level based on a series of developmentally-orchestrated mitochondrial processes. We demonstrate that mitochondrial fission, together with the lack of mtDNA replication in early germarium, effectively segregates mtDNA into individual organelles. After mtDNA segregation, mtDNA transcription begins, which leads to the activation of respiration in each organelle. The expression of mtDNA-encoded genes allows the functional manifestation of different mitochondrial genotypes in heteroplasmic cells, and hence functions as a stress test for each individual genome and sets the stage for the replication competition. We also show that the Balbiani body has a minor role in mtDNA selective inheritance by supplying healthy mitochondria to the pole plasm. The two selection mechanisms may act synergistically to secure the transmission of functional mtDNA through *Drosophila* oogenesis.

## Introduction

Mitochondria, the indispensable power plants of eukaryotic cells, present geneticists with a fundamental paradox. Their genome accumulates mutations at a high rate in the soma, estimated to be two orders of magnitude higher than that of the nuclear genome (Wallace and Chalkia, 2013). This rate owes to both the abundance of highly-mutagenic free radicals generated by respiration and the lack of effective DNA repair *via* homologous recombination in the mitochondrial matrix (Taylor and Turnbull, 2005). If freely transmitted, the damaging mutations would gradually accumulate over generations, which could severely impair the fitness of organisms and even lead to their extinction (Felsenstein, 1974). However, damaging mtDNA mutations are exceedingly rare in populations. This paradox underscores the existence of effective mechanisms that restrict the transmission of deleterious mtDNA mutations and favor the selective inheritance of healthy mitochondrial genomes. Since mtDNA is predominantly transmitted through the maternal lineage, these mechanisms must operate in the female germline.

Currently, the dominant dogma, bottleneck inheritance proposes that only a small fraction of the mitochondrial genomes present in primordial germ cells are transmitted to an oocyte and eventually populate the offspring. This process explains the rapid genetic drift of mitochondrial genotypes between generations (Hauswirth and Laipis, 1982; Olivo et al., 1983; Rebolledo-Jaramillo et al., 2014). Bottleneck inheritance could also indirectly lead to the counterselection of deleterious mtDNA mutations: different mtDNA compositions would generate different metabolic outputs in developing oocytes, eventually causing the elimination of oocytes harboring an excess of deleterious mutations (Jenuth et al., 1996). However, the frequency of spontaneous mtDNA mutations is about 10^−5^−10^−6^ (Cree et al., 2008), which, while high compared to the frequency of nuclear DNA mutations, remains too low to elicit the kind of biochemical deficiency required for an effective selection at the whole-cell level. In fact, deleterious mtDNA mutations are prevented from passing to the next generation even when present in low copy-number in mouse models (Fan et al., 2008; Stewart et al., 2008b), underscoring the existence of a selection that likely occurs at the level of individual organelles or genomes, besides the model of bottleneck inheritance.

Our work in *Drosophila melanogaster* (*Dm*) has shown that selective inheritance of mtDNA also takes place in insects (Hill et al., 2014). Taking advantage of a temperature-sensitive deleterious mtDNA mutation (*mt:CoI*^*T300I*^, referred to as *ts*) that we had previously engineered (Hill et al., 2014), we found that the load of *ts* allele in the progeny of heteroplasmic mothers (carrying both wild-type and *ts* mtDNAs) was greatly reduced at restrictive temperature (Hill et al., 2014). This observation suggested that the *Dm* female germline could detect the defect caused by the *ts* allele at restrictive temperature, and limit its transmission. Genetic and developmental analyses roughly mapped selective mtDNA inheritance in *Dm* to a developmental window spanning the late germarium (region 2B) and early stages of egg chamber (Hill et al., 2014; Ma et al., 2014). This is a stage where groups of 16 sister cells (16-cell cysts), derived from the four successive divisions of single oocyte precursors, organize into egg chambers from which a single egg will emerge (Spradling, 1993). Interestingly, this is also the stage when mtDNA replication resumes, after having been largely quiescent in the earlier dividing cysts and region 2A (Hill et al., 2014). mtDNA replication in this region appears to depend on active mitochondrial respiration, as both pharmacological inhibition and genetic disruption of nuclear-encoded electron-transport chain (ETC) subunits severely impair mtDNA replication (Hill et al., 2014). The *ts* mutation, which disrupts a subunit of the mitochondria-encoded ETC complex IV, also leads to greatly diminished mtDNA replication in homoplasmic germaria that carry only the mtDNA *ts* allele under restrictive temperature (Hill et al., 2014). We reasoned that in heteroplasmic germ cells, healthy mitochondria carrying wild-type mtDNA would replicate their DNA and propagate much more vigorously than defective ones harboring mutations that impair respiration, which would consequently reduce the proportion of mutant mtDNA in the progeny. This reasoning led us to propose a replication-competition model for selective inheritance in *Dm* (Hill et al., 2014).

While logically compelling, the replication-competition model rests on the assumption that germ cells can discern the integrity of individual mitochondrial genomes, presumably based on their functionally distinct protein products. However, mitochondria are not stationary: they undergo constant fusion and fission, which mixes and reassorts mitochondrial genomes and their products (Frank et al., 2001; Ishihara et al., 2006; Legros et al., 2004). In heteroplasmic cells, mitochondrial fusion allows mtDNA complementation to maintain the overall metabolic output in spite of significant levels of mtDNA mutations (Chan, 2006). This process could therefore mask the functional deficiency caused by deleterious mutations, and prevent the elimination of defective genomes. Therefore, we propose that mitochondrial genomes have to be effectively segregated, and then expressed, for replication-competition to be effective in selective mtDNA inheritance. However, currently, it is not clear whether and how mtDNA segregation and expression are regulated during oogenesis, nor do we know how such regulation impacts mtDNA transmission and selective inheritance.

Aside from replication-competition and bottleneck inheritance, another mechanism has also been proposed, based on the observed localization and transport of mitochondria to the prospective germ plasm during oogenesis (Cox and Spradling, 2003). At the beginning of oocyte differentiation, a fraction of mitochondria and other organelles congregate within a structure called the Balbiani body, which supplies mitochondria to the pole plasm of the mature oocyte, the cytoplasm of the future embryo’s primordial germ cells. It has been proposed that healthy mitochondria might be enriched in the Balbiani body and preferentially transmitted to grandchildren (Cox and Spradling, 2003). However, this idea still lacks direct experimental support. Since we have shown that selective inheritance was detectable as early as in the offspring of heteroplasmic mothers, not just in their grandchildren, this mechanism could not alone explain selective inheritance. Nevertheless, it might be an important contributor.

In this paper, we test the basic assumptions of our replication-competition model by documenting the behavior of mitochondria and their genomes in the developing *Dm* ovary. In addition, we examine the potential contribution of mitochondrial aggregation around the Balbiani body to selective mtDNA inheritance in *Dm*.

## Results

### Mitochondrial morphological change in *Dm* early germarium

Using optical microscopy, previous studies have demonstrated that mitochondria were more rounded and fragmented in dividing cysts compared to germline stem cells (GSCs) in *Dm* germarium region 1 (Cox and Spradling, 2003; Lieber et al., 2019), suggesting that mitochondria undergo increased fission or decreased fusion in germline cysts. To comprehensively analyze mitochondrial size and shape in germarium, we used focused ion beam scanning electron microscopy (FIB-SEM) to reconstruct a 3D volume of *Drosophila* germarium at an isotropic resolution of 10×10×10 nm^3^ voxels (Videos, 1 and 2). We then applied the computational segmentation to trace all mitochondria in germ cells. Overall, mitochondria displayed a wide spectrum of morphology, ranging from small spheres and short tubules to elongated tubules in germarium (Fig. 1A and B). The mitochondrial volume appeared to be gradually decreased during cyst division (Fig. S1 A). In a 2-cell cyst at region 1, 47.9% of the mitochondria were larger than 0.05 µm^3^ (Fig. 1 E). After the completion of cyst division, the fraction of large mitochondria (>0.05 µm^3^) decreased to 18.8% in a 16-cell cyst at region 2A (Fig. 1 E). Reciprocally, the fraction of small mitochondria (<0.03 µm^3^) increased to 63.5%, compared with 37.5% in region 1 (Fig. 1 C, Table S1). We plotted the volume against the surface area of each individual mitochondrion to evaluate the geometric shape of mitochondria (Harris and Theriot, 2018) (Fig. S1 B and C). In region 1, two populations of mitochondria were observed (Fig. S1 B). One group of mitochondria were spherical, while others were more elongated. In region 2A, more mitochondria shifted toward spheroid morphology (Fig. S1 C). These observations, taken together, suggest that elongated, large mitochondria undergo fragmentation in diving cysts in region 1, and become smaller spheroids in 16-cell cyst region 2A.

**Figure 1.**
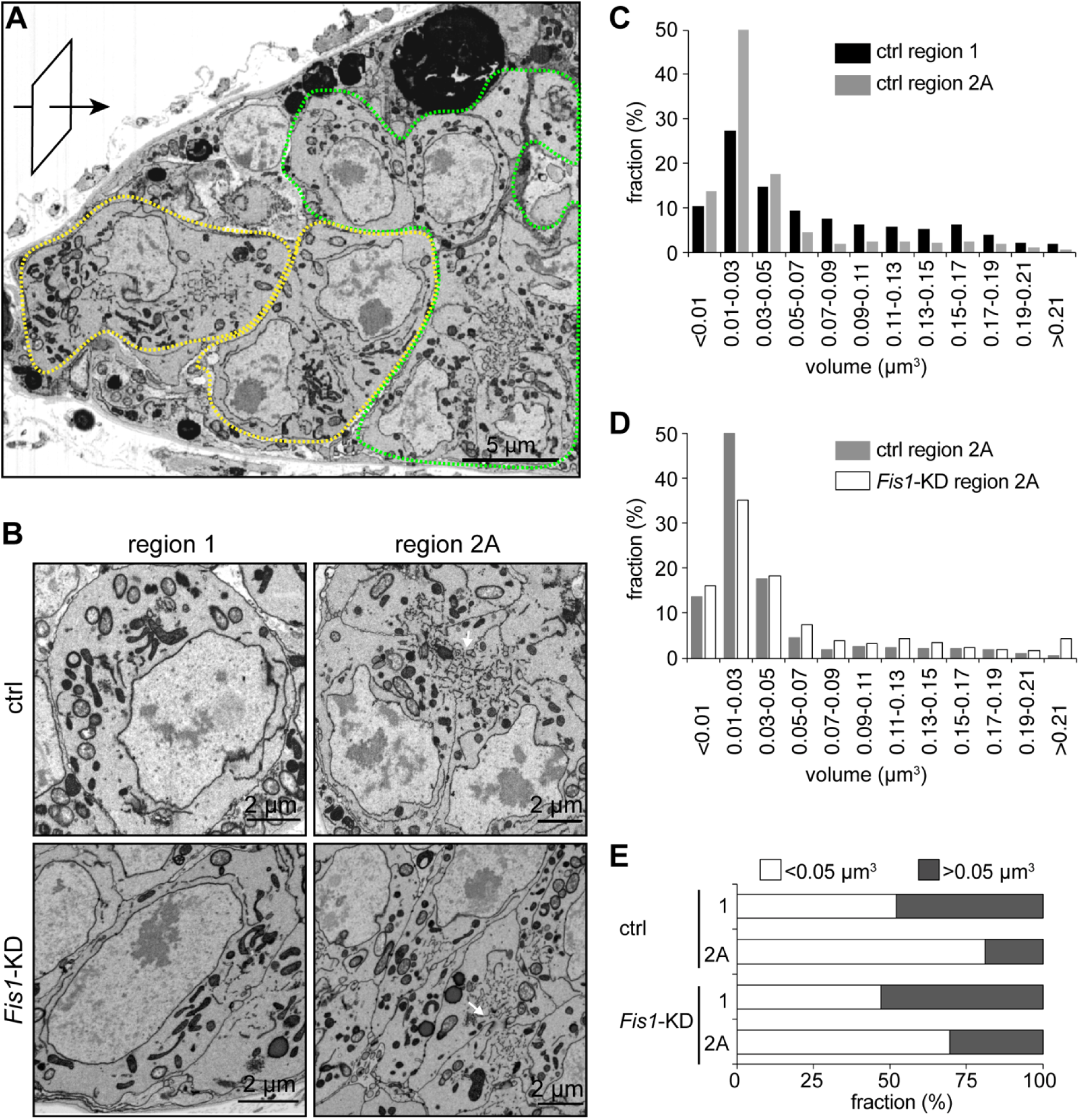
FIB-SEM show mitochondrial fragmentation in *Dm* early germarium, which could be compromised by *Fis1* knockdown. **(A)** A representative electron micrograph of wild-type early germarium obtained by FIB-SEM. Sample milling was performed from the anterior tip of the germarium towards posterior (arrow). The image is a single section of xz plane after 3D reconstruction. Region 1 and region 2A cysts are outlined with yellow and green dashed lines, respectively. Scale bar, 5 µm. **(B)** The detailed structures of mitochondria and other subcellular components in region 1 and region 2A of wild-type and *Fis1* knockdown by *bam-gal4*. The arrows point to the ring canals between the cystocytes and the fusome that extends through the ring canals, which were used to trace the germ cells within a cyst. Scale bar, 2 ⍰m. **(C and D)** Frequency distribution of mitochondrial volume in region 1 and region 2A of wild-type germarium **(C)**, and those in region 2A of wild-type and *Fis1* knockdown germarium **(D)**. The proportion of mitochondria fall to the indicated range of volume are shown. **(E)** The proportion of mitochondria with bigger (> 0.05 µm3) volume from each experimental group.

To confirm this observation, we assessed mitochondrial morphology in germarium expressing dsRNA against *Fis1*, a mitochondrial outer membrane protein that promotes mitochondria fission (Stojanovski et al., 2004). *Fis1* RNAi was activated by a *bam-gal4* driver that expresses in dividing cysts specifically (Chen and McKearin, 2003) (Fig. S2 A). There were more large mitochondria (>0.05 µm^3^), and correspondingly, less small mitochondria (0.01-0.03 µm^3^) in region 2A compared to the control (Fig. 1 D). The results indicate that *Fis1* knockdown can effectively impede mitochondrial fragmentation in early germarium.

### Mitochondrial genomes are segregated in region 2A of the *Drosophila* germarium

We reasoned that the lack of mtDNA replication in region 2A (Hill et al., 2014), in conjunction with mitochondria fragmentation would facilitate mitochondria genome segregation. To test this idea, we visualized mtDNA and their distribution in the mitochondrial network in germarium. Mitochondrial transcription factor A (TFAM) is the major mtDNA packaging protein and a well-established marker for mtDNA nucleoids (Alam et al., 2003). We thus used stimulated emission depletion (STED) microscopy to image both mitochondria and TFAM-GFP (Zhang et al., 2016), to assess mtDNA segregation (Fig. 2 A). Mitochondria that were labelled by ATP synthase α subunit (ATPs) staining were more elongated and interconnected in stem cells or cystoblasts at region 1, whereas became more rounded in region 2A (Fig. 2 A and B). This result further substantiates that mitochondria undergo fragmentation in early germarium as observed in the FIB-SEM analysis. The TFAM-GFP signal showed as many puncta throughout the germarium and localized to mitochondria. We noticed that the ATPs staining was not always uniform along the mitochondrial network, which might reflect the uneven density of cristae, where the ATP synthase locates, in different mitochondria. Sometimes, even within a single mitochondrion, ATPs staining appeared as multiple puncta, and nucleoids were located in regions with low ATPs intensity (Fig. 2 B). This phenomenon is in line with a recent super-resolution microscopic study showing a lack of cristae structures surrounding nucleoids (Stephan et al., 2019).

**Figure 2.**
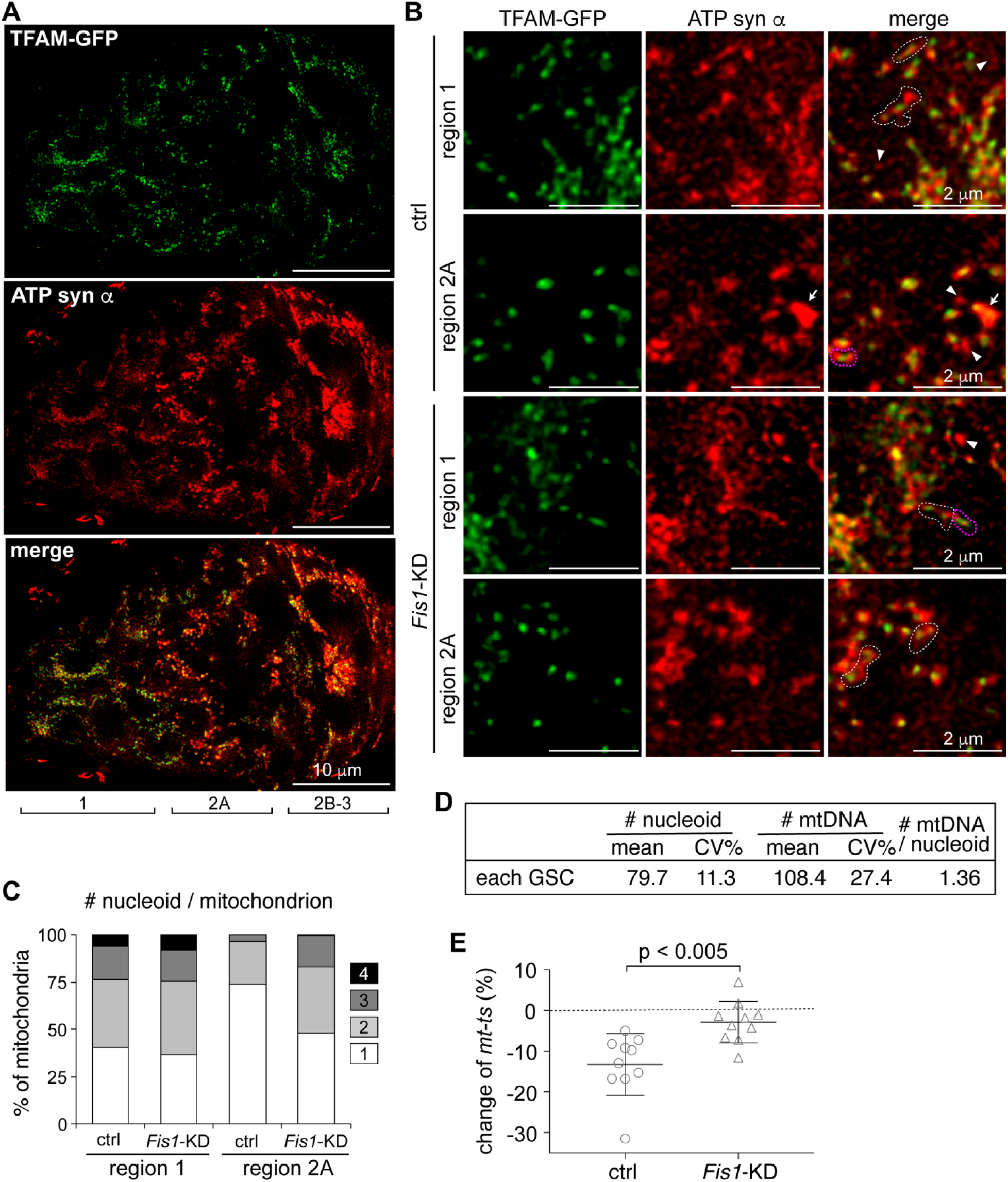
Mitochondrial fragmentation and nucleoid segregation in germarium region 2A are essential for mtDNA selective inheritance. **(A)** Mitochondria labeled by ATP synthase ⍰ subunit staining and mtDNA labeled by TFAM-GFP in *Drosophila* germarium. The images are representative *z*-stack projection (1.5 µm) of a germarium using stimulated emission depletion (STED) microscopy. The developmental regions of germarium are indicated. Images here and throughout are oriented with the germarium anterior towards the left. Scale bar, 10 µm. **(B)** Mitochondrial nucleoids labeled by TFAM-GFP and mitochondria labeled by ATP synthase ⍰-subunit staining in a germarium are displayed in magnified views. Both control and *Fis1* knockdown ovaries driven by *bam-gal4* are shown. The representative images of region 1 are from anterior end, where germline stem cells or cystoblasts reside. The white dashed outlines are elongated mitochondria containing multiple nucleoids. Arrowheads point to mitochondria without nucleoids. The arrow point to the clustered mitochondria, which are excluded from data analyses. The ATP synthase α subunit staining were uneven, with less intensity in the regions where nucleoids are located. We defined two adjacent, but distinct ATP synthase subunit α puncta as a single mitochondrion if they appeared in the same contour and were connected with one nucleoid (magenta dashed outlines). Scale bar, 2 ⍰m. **(C)** The number of nucleoids per mitochondrion was determined using TFAM-GFP and ATP synthase ⍰-subunit staining shown in **(B)** in regions 1 (stem cells or cystoblasts) and 2A from control and *Fis1* knockdown ovaries (n = 10 cysts for each group). The fraction of each group is shown. Note that the fraction of mitochondria containing multiple nucleoids was increased in region 2A of *Fis1* knockdown driven by *bam-gal4*. **(D)** Quantification of mitochondrial nucleoid number and mtDNA copy number in each germ stem cell (GSC) in germarium. CV, coefficient of variance; **(E)** Knockdown of *Fis1* in germarium region 2A, using a *bam-Gal4* driver, compromises the selection against the mutant mtDNA in heteroplasmic *mt:CoI^T300I^ Drosophila*. The proportion of mutant *ts* mtDNA in progeny is decreased by 15% compared with their mothers on average. In *Fis1* knockdown fly, this negative selection was diminished.

We next quantified the number of nucleoids in each mitochondrion. In the anterior end of region 1, where stem cells and cystoblasts reside, the number of mitochondrial nucleoids, indicated as the TFAM-GFP puncta, ranged from zero to four in different mitochondria (Fig. 2 C). As expected, large and elongated mitochondria often contained more than one nucleoids, whereas some small mitochondria that might be the intermediate structures of mitochondrial fusion and fission processes, were devoid of TFAM-GFP signal (Fig. 2 B). Among TFAM-GFP positive mitochondria, 40.4%, 36% 17.4% and 6.2% of them contained 1, 2, 3 and 4 nucleoids, respectively (Fig. 2 C). In region 2A, 73.9% of the mitochondria contained only 1 nucleoid (Fig. 2 C), indicating that through the mitochondrial fragmentation, nucleoids are effectively segregated in 16-cell cysts at region 2A before the onset of mtDNA replication in germarium region 2B.

It has been estimated that a single nucleoid may contain 1 to 10 copies of mtDNA (Kukat et al., 2011; Legros et al., 2004; Satoh and Kuroiwa, 1991). Although mtDNA in separate nucleoids do not intermix (Gilkerson et al., 2008), those within a nucleoid can functionally complement each other, which could interfere with selective inheritance (Hill et al., 2014). Currently, there is no reliable technique to accurately quantify mtDNA copy number within a specific nucleoid. We considered using TFAM-GFP intensity as a measure for mtDNA copy number in a nucleoid. However the values of TFAM-GFP intensities in different nucleoids appeared to be random and continuous, instead of quantized. Therefore, the TFAM-GFP intensity is not only determined by the mtDNA copy number, but could also affected by the compaction state of mtDNA (Kukat et al., 2011). Nonetheless, one could approximate mtDNA copy number per nucleoid by normalizing the total number of mitochondrial genomes to the number of nucleoids in a cell. To this end, we isolated germ cells by Fluorescence Activated Cell Sorting from the ovary of white pupae expressing Vasa-GFP, a reporter specific to germ cells (Fig. S3 A). At this stage, oogenesis has progressed to region 1 of the germarium, which consists mainly of germline stem cells (GSCs) and cystoblasts (Song et al., 2007). We quantified the mtDNA copy number to be ∼108 copies in GSCs and early cysts (Figure 2D). We also quantified the mtDNA copy number in a female germline stem cell culture (fGS) established from *Drosophila* adult ovaries (Niki et al., 2006). We estimated there were about 120 copies of mtDNA per fGS cell (Fig. S3 B), which is close to the value obtained from germ cells in early pupae ovaries. Since there were approximately 80 nucleoids in GSCs (Fig. 2 D), we deduced that each nucleoid contained 1.36 copies of mtDNA on average. This number suggests that intra-nucleoid complementation is rather minimal at this stage. In addition, given that there is no mtDNA replication until region 2B (Hill et al., 2014), the mtDNA copy number within a nucleoid should not increase prior to the stage when selective inheritance occurs. Taken together, our observations indicate that mtDNA molecules are effectively sorted into different organelles during the early stages of ovarian development.

### Mitochondrial fragmentation in region 2A is required for mtDNA selective inheritance

To test whether mtDNA segregation was required for selective inheritance, we attempted to increase the number of nucleoids per mitochondrion by tampering with mitochondrial fission. We quantified the nucleoid numbers in individual mitochondrion in *Fis1* knockdown flies (Fig. 2 B and C). In 16-cell cyst region 2A, only 47.7% of mitochondria contained a single nucleoid, compared to 73.9% in wild-type (Fig. 2 C). Additionally, RNAi against *Drp1* (Fig. S4 A and B), a small GTPase that promotes mitochondria fission (Labrousse et al., 1999), caused similar phenotypes as these in *Fis1* knockdown flies. Thus, inhibition of mitochondrial fission impairs mitochondria genome segregation.

We next tested the impact of impaired mitochondrial fission on mtDNA selective inheritance in heteroplasmic flies. We knocked down *Fis1* or *Drp1* in the female germline of the heteroplasmic fly using *bam-gal4* driver and quantified heteroplasmy in mothers and their eggs. Under the restrictive temperature, the average proportion of *ts* allele was decreased by 15% in eggs compared to their mothers in control flies (Fig. 2 E), indicating a selection against the deleterious mtDNA. By contrast, in both *Fis1* RNAi and *Drp1* RNAi flies, the load of *ts* allele in progeny displayed a pattern of random mtDNA segregation, with no clear decrease in the proportion of *ts* genomes (Fig. 2 E; Fig. S4 C). These results suggest that the inhibition of mitochondrial fission at region 2A weakened mtDNA selective transmission. Taken together, these observations indicate that mitochondrial fission promotes nucleoid segregation in region 2A and is required for effective mtDNA selection.

### Mitochondrial activity and mitochondrial DNA expression commence in region 2B

We hypothesize that for heteroplasmic cells to distinguish between mitochondria with different mtDNA genotypes, ETCs genes on mitochondrial genomes have to be expressed to assess their functional readout: mitochondrial respiration. To test this idea, we examined the pattern of mtDNA-encoded gene expression in the germanium by fluorescence *in situ* hybridization (FISH), using short DNA probes targeted to two mtDNA encoded mRNAs: *NADH dehydrogenase 4 (ND4)* and *cytochrome c oxidase subunit 1 (cox1)* (Fig. 3 A). For either *ND4* or *cox1*, moderate level of mRNA was detected in GSCs, but almost no signal in region 2A. In the following 16-cell cyst at region 2B, a strong mRNA signal was observed, suggesting that mtDNA expression commences at this stage (Fig. 3 B). We also checked the expression pattern of two ETC subunits encoded in the nuclear genome: *NADH: ubiquinone oxidoreductase subunit B5 (NDUFB5)* and *cytochrome c oxidase subunit 5A (cox5A)* (Fig. 3 A). Both transcripts showed an expression pattern similar to that of mtDNA-encoded mRNAs, i.e., moderate expression in germline stem cells, nearly no expression in region 2A, and strong expression in region 2B (Fig. 3 B). Interestingly, the localization of these two transcripts resembled that of mitochondria at the region 2B germarium, which is consistent with previous studies that mitochondrial ETCs genes undergo local translation on mitochondrial outer membrane (Williams et al., 2014; Zhang et al., 2016). The coordinated expression of mitochondrial and nuclear genes encoding subunits of the ETC in region 2B led us to ask whether mitochondrial respiration was also activated at this stage.

**Figure 3.**
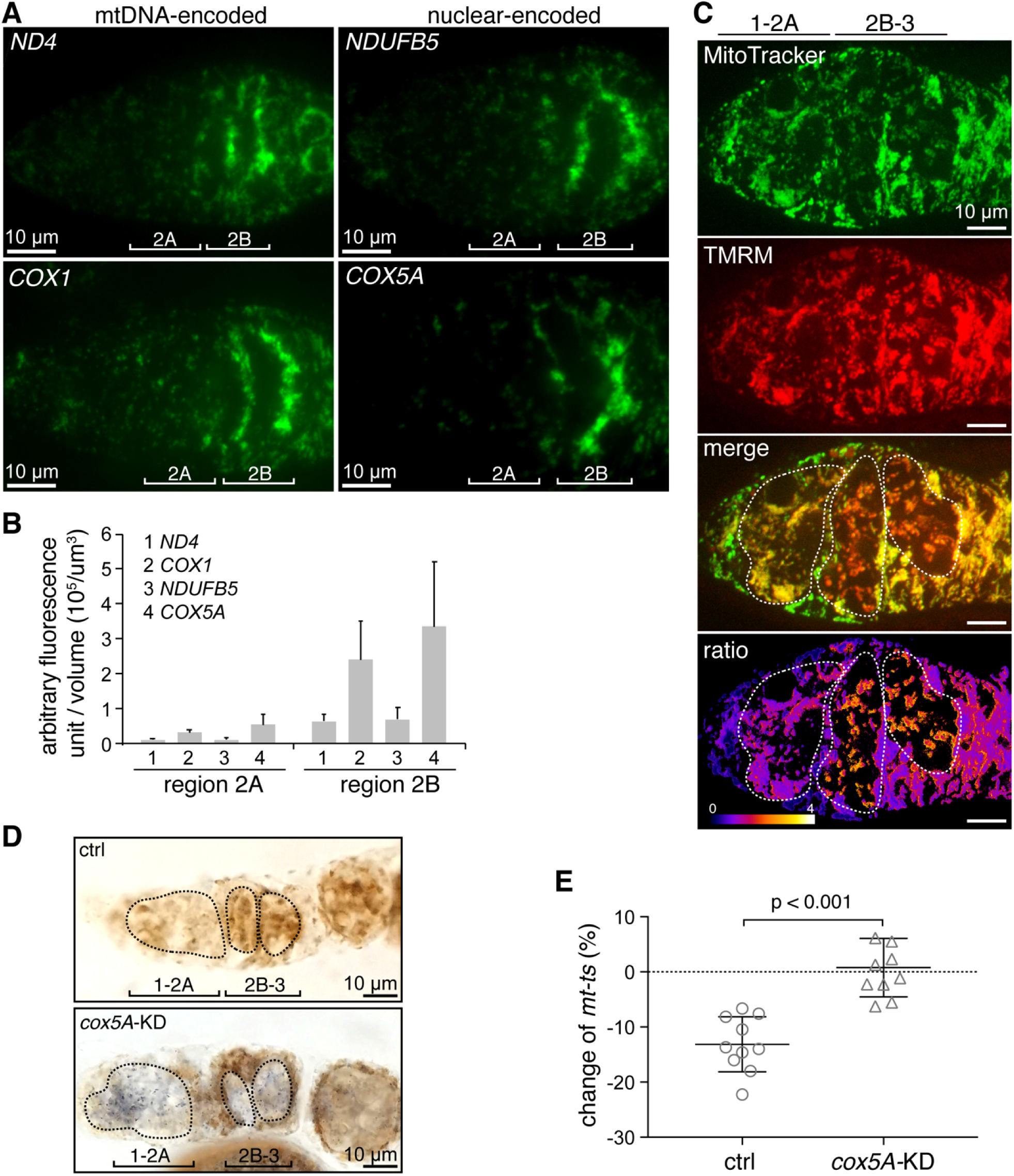
Mitochondrial respiration is activated in region 2B and essential for mtDNA selective inheritance. **(A)** Expression of mtDNA- and nuclear-encoded ETCs genes in the germarium. The spatial patterns of nuclear and mitochondrial encoded mRNAs were revealed by smFISH assay. Fluorescently labelled probes targeted mtDNA-encoded *ND4* and *cox1* transcripts, and the nuclear-encoded *NDUFB5* and *cox5A* transcripts. Scale bar, 10 µm. **(B)** Quantification of total immunofluorescence intensity per volume in region 2A and 2B. The expression of both types of RNAs were markedly increased at region 2B. **(C)** Mitochondria membrane potential staining demonstrated by the mitochondrial membrane potential indicator TMRM (red) and mitochondrial fluorescent dye mitoTracker (green). The strong red signal at region 2B and 3 in merged image suggests markedly increased membrane potential. The ratio of red to green fluorescence intensity is shown as the pseudo color ratiometric image. The developing regions of germarium germ cells are outlined. Scale bar, 10 µm. **(D)** Mitochondrial respiratory activity in ovary using the colorimetric assay. Ovaries from control and *cox5A* knockdown flies driven by *nanos-gal4* were stained for dual succinate dehydrogenase (complex II) and cytochrome C oxidase (complex IV) activity. Representative images for each group are shown. Intense brown color, indicating that both complex II and complex IV are active, was prominent in region 2B, but mostly absent in region 1 and region 2A. The strong blue color in *cox5A* knockdown group suggests complex IV activity was greatly reduced upon *cox5A* knockdown at all developmental stages of germarium. The developing regions of germarium germ cells are outlined. Scale bar, 10 µm. **(E)** Selection against the deleterious mtDNA mutation (*ts*) in heteroplasmic *mt:CoI*^*T300I*^ *Drosophila* was compromised by knocking down *cox5A* in the germline using *nanos-gal4* driver. *p*<0.001.

To address this question, we analyzed mitochondrial membrane potential during germarium development, using tetramethylrhodamine methyl ester (TMRM), a dye that accumulates in polarized mitochondria (Scaduto and Grotyohann, 1999). We found that the ratio of TMRM to MitoTracker green (a marker of mitochondrial mass) was markedly increased in germanium region 2B compared to regions 1 and 2A (Fig. 3 C), indicating that mitochondrial membrane potential is low in early-stage cysts, but up-regulated at the region 2B. The low level of membrane potential in the early germarium might simply owe to a lack of ETCs. To assess the level of ETCs, we directly evaluated activities of ETCs in a colorimetric assay by incubating ovaries with substrates of succinate dehydrogenase (complex II) and cytochrome C oxidase (complex IV) (Ross, 2011). Intense brown color, indicating that both complex II and complex IV are present and active, was evident in region 2B, but mostly absent in region 1 and region 2A (Fig. 3 D). This result is consistent with the mitochondrial membrane potential staining (Fig. 3 C), indicating that mitochondrial respiration is low in early germarium, but elevated in region 2B. Given their co-occurrence, the elevation of respiration is at least, partially due to the onset of mtDNA expression that generates ETCs in differentiating cysts region 2B, when selective inheritance begins.

### Mitochondrial activation in late germarium stage allows the selective propagation of functional mtDNA

Based on the results above, we hypothesized that mtDNA expression and the following activation of respiration may act as a stress test allowing germ cells to distinguish between mitochondria that harbor a wild-type versus a mutant mtDNA. We hence predicted that a ubiquitous disruption of mitochondrial activity in a heteroplasmic germ cell, would mask the deficiency of mitochondria harboring the deleterious mtDNA mutation, and impair selective inheritance. To disrupt respiration in all mitochondria, we knocked down a nuclear-encoded ETC gene, *cytochrome c oxidase subunit 5A* (*cox5A*) in ovaries driven by *nanos-gal4* (Fig. S2 A). Strong knockdown of *cox5A* led to degeneration of ovaries. We were able to find an appropriate RNAi line that moderately decreased the mRNA level of *cox5A* (Fig. S2 D and F), but did not affect the fecundity of the female flies or the hatching rate of their progeny (Fig. S5). Nonetheless, complex IV activity was markedly decreased in the knockdown germarium (Fig. 3 D). Importantly, knockdown of *cox5A* in heteroplasmic *mt:CoI*^*T300I*^ background severely impaired the selection against the *ts* allele (Fig. 3 E).

The universal disruption of mitochondrial respiration by knocking down *cox5A* in a germ cell, not only masks the differential energetic status among different mitochondria, but also could potentially impair cellular energy metabolism. If the activation of mitochondrial respiration in region 2B indeed functions as the stress test for mtDNA integrity, we anticipated that improving the respiratory activity of defective mitochondria carrying deleterious mutations would also weaken selective inheritance in heteroplasmic flies. We previously showed that the ectopic expression of an alternative oxidase, AOX, that catalyzes electron transfer from ubiquinone to molecular oxygen and bypasses the cytochrome chain reactions, completely restored the viability of *mt:CoI*^*T300I*^ flies (Chen et al., 2015). Expression of AOX in the germ cells driven by *nanos-gal4* partially restored mtDNA replication in region 2B of homoplasmic *ts* flies raised at 29 °C, based on an EdU incorporation assay (Fig.4 A and C). Since mtDNA replication in region 2B depends on active respiration (Hill et al., 2014), this observation confirms that AOX overexpression rescues the respiration defect of *ts* mitochondria at restrictive temperature. When we overexpressed AOX in heteroplasmic flies, we found the reduction of *ts* allele in progeny was much less pronounced than in control flies (Fig. 4 D). This observation indicates that mitigating the mitochondrial deficiency caused by *ts* mtDNA impairs the selection process.

**Figure 4.**
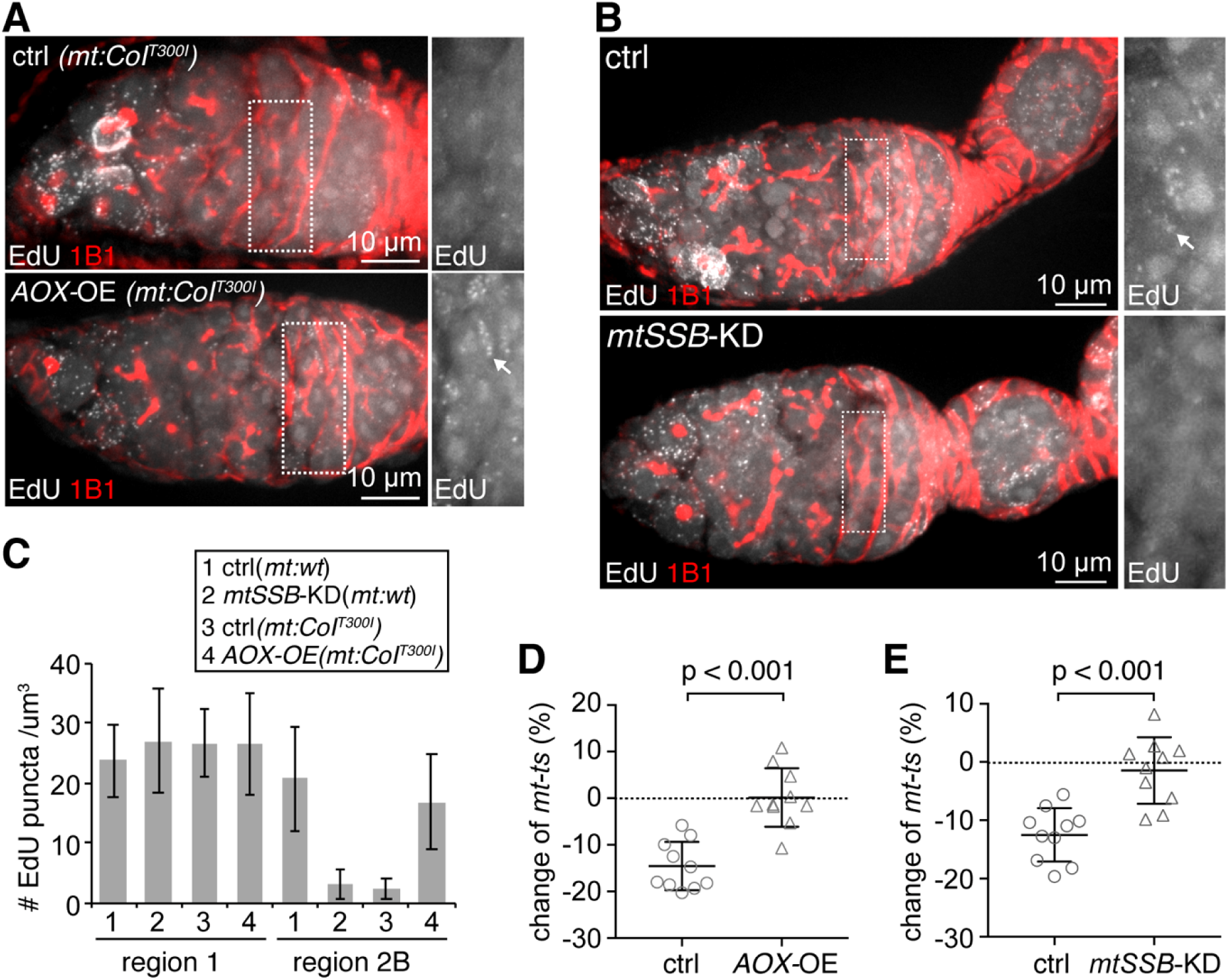
mtDNA replication are indispensable for selective inheritance. **(A and B)** mtDNA replication labeled by EdU staining in *Drosophila* germarium. The Hts-1B1 antibody was used to stain the fusome, which indicates the developmental stages of germarium. Region 2B is outlined in white and enlarged on the right panels. Note that the mtDNA replication was specifically disrupted at region 2B in *mt:CoI*^*T300I*^ ovary at 29°C, but could be restored by overexpression of AOX driven by *nanos-gal4* **(A).** Knocking down mtSSB using *nanos-gal4* driver diminished mtDNA replication at region 2B of wild-type ovary, but not in egg chambers **(B).** Arrows points to the EdU puncta. Scale bar, 10 ⍰m. **(C)** Quantification of mtDNA replication indicated by numbers of EdU puncta in regions 1 and 2B in genetic backgrounds shown in **(A)** and **(B). (D and E)** Selection against *ts* mtDNA at 29°C was compromised by ectopic expression of AOX **(D)** and knock down of *mtSSB* **(E).** Both lines were driven by *nanos-gal4. p*<0.001.

The fact that improving respiration in heteroplasmic flies without affecting the overall cellular energy metabolism, can weaken selective inheritance suggests that germ cells indeed rely on the respiratory activity of individual mitochondria to gauge the integrity of their mtDNA. Based on our replication-competition model, respiratory activity would then promote selective mtDNA inheritance by allowing the wild-type mtDNA harbored by healthy mitochondria to replicate more efficiently than the mutant mtDNA harbored by respiration-defective organelles. This model predicts that mtDNA replication would be required for selective inheritance. To test this idea, we attempted to inhibit mtDNA replication in heteroplasmic flies. We found that knockdown mitochondrial single-stranded DNA binding protein (*mtSSB*) (Fig. S2 E), an essential factor for mtDNA replication (Korhonen et al., 2004), markedly reduced mtDNA replication in region 2B, while mtDNA replication in later egg chambers appeared unaffected (Fig. 4 B). This result is consistent with the notion that mtDNA replication in region 2B is particularly sensitive to mitochondrial disruption (Hill et al., 2014). Importantly, knocking down *mtSSB* in heteroplasmic flies greatly diminished selective inheritance (Fig. 4 E). Additionally, RNAi against *tamas*, the mitochondrial DNA polymerase also diminished the selection against the *ts* mtDNA (Fig. S 6), supporting that mtDNA replication in region 2B is indeed necessary for selective inheritance. Taken together, the results described above suggest that activation of mitochondrial respiration serves as a stress test that identifies healthy mitochondria and promotes the replication of their mtDNA.

### Balbiani body makes a small contribution to selective inheritance in germ cells

So far, we have shown that mtDNA molecules are effectively segregated before region 2B and begin expressing their genes and replicating in region 2B. In a heteroplasmic background, these concerted behaviors presumably allow the healthy mitochondria containing wild-type mtDNA to outcompete the defective mitochondria harboring deleterious mutations in developing germ cells, which effectively reduces the proportion of mtDNA mutations in mature oocytes.

However, a developmentally-regulated localization and transport of mitochondria in the germarium has also been proposed to contribute to mtDNA selection (Cox and Spradling, 2003). In germarium region 2B, healthy mitochondria are preferentially associated with fusome (Hill et al., 2014). Some fusome-associated mitochondria will be transported to the Balbiani body and populate in the pole plasm (COX and Spradling 2006), the cytoplasm of future primordial germ cells (PGCs). Thus, one would expect the PGCs of an embryo to have a lower level of mtDNA mutations than its somatic cells. To test this idea, we isolated PGCs from the fertilized eggs of heteroplasmic flies expressing a germ cell specific reporter, Vasa-GFP, by fluorescence-based cell sorting, and compared levels of the *ts* allele in PGCs and somatic cells. At 29 °C, the proportion of *ts* genomes in PGCs was slightly, but consistently, lower than in somatic cells in the same batch of embryos (Figure 5A, ctrl). The difference in heteroplasmic level between PGCs and somatic tissues was about 3% on average. To confirm that this difference results from an enrichment for healthy mitochondria in the Balbiani body, we performed the same experiment in the context of a germline-specific knockdown of *milton*, the adaptor that mediates the Kinesin-dependent transport of mitochondria to the Balbiani body (Cox and Spradling, 2006). In forming follicle, there were much less mitochondria at the anterior end of the *milton* knockdown oocyte compared to the control (Fig. 5 B). Mitochondria dispersed throughout the cytoplasm of the oocyte in the control stage 5 egg chambers. However, *milton* knockdown oocytes contained much fewer mitochondria, most of which remained at the anterior end (Fig. 5 B). These phenotypes resemble those of a *milton* null mutant (Cox and Spradling, 2006), indicating the effective disruption of both Milton activity and Balbiani body formation. Of primary importance, the load of *ts* allele was similar between PGCs and somatic cells in *milton* knockdown flies (Fig. 5 A), suggesting that a Balbiani body-associated selection of mitochondria does take place. This selection plays a complement role to further improve mitochondrial fitness in germ cells and may act synergistically with other mechanisms to further enhance the selective transmission of mtDNA in *Dm*.

**Figure 5.**
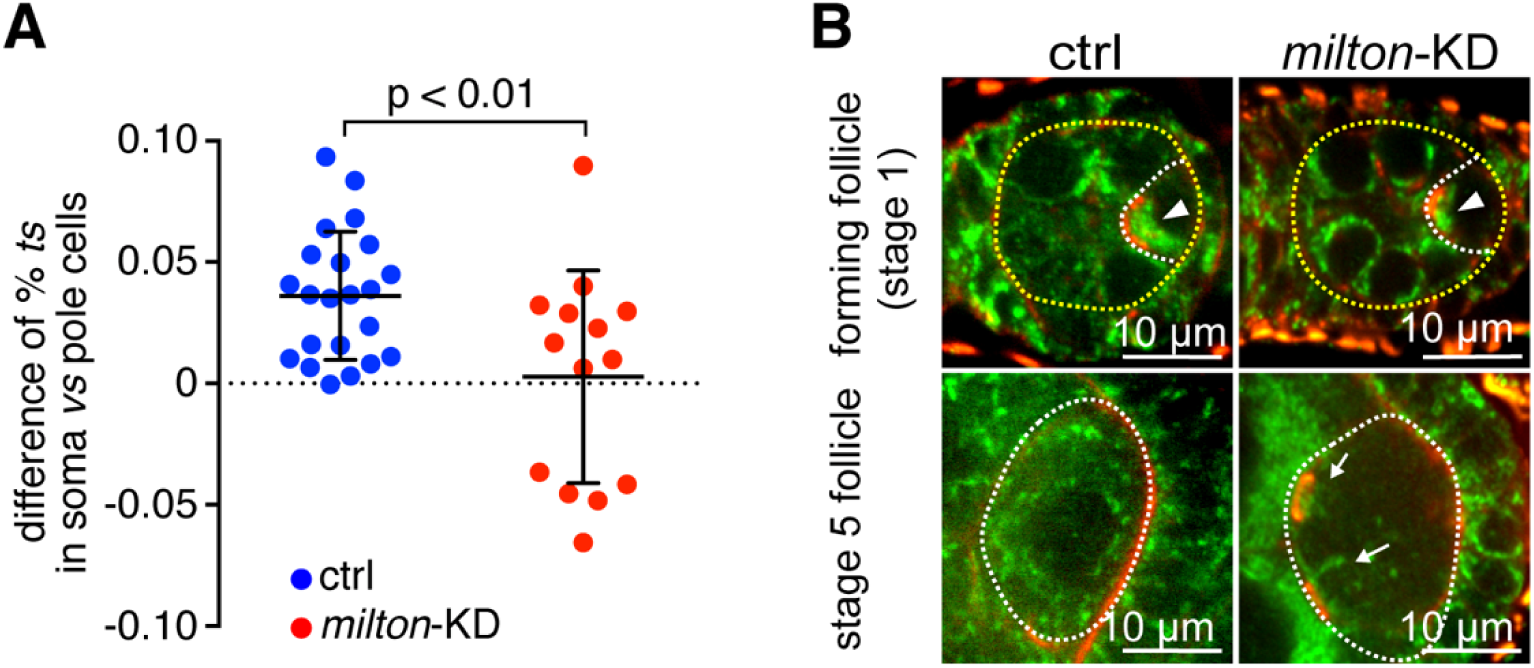
Balbiani body contributes to the selection against the *mt:CoI*^*T300I*^ mtDNA in the germline of heteroplasmic flies. **(A)** Somatic cells consistently had a higher percentage of *ts* mtDNA than germ cells (pole cells) in heteroplasmic embryos. Disrupting the Balbiani body by knocking down *milton* gene (*milton-KD*) in germ cells abolished the difference between somatic and germ cells. **(B)** Balbiani body formation was disrupted in *milton* knockdown oocytes. Mitochondria labeled by ATP synthase α subunit staining (green), ring canals and other actin filaments stained by phalloindin (red) are shown. In the oocyte (white dashed outlines) of the forming follicle cyst (yellow dashed outlines), mitochondria in *milton* knockdown were less and remained at the anterior of the oocyte. A normal Balbiani body (arrowheads) could not be formed. In the oocyte (white dashed outlines) of stage 5 wild-type follicle, mitochondria were sparse and distributed evenly in cytoplasm. There were much fewer mitochondria, which located near ring canals (arrows) in *milton* knockdown oocyte. Scale bar, 10 µm.

## Discussion

In this study, we illustrate a series of developmentally-orchestrated mitochondrial processes in *Dm* germarium, including mitochondrial fragmentation in early germarium cysts, mtDNA expression and ETCs activation in late germarium region 2B, that are indispensable for restricting the transmission of deleterious mtDNA mutations (Fig. 6). Using FIB-SEM and computational segmentation, we comprehensively documented mitochondrial morphology in *Drosophila* germarium. We found that mitochondria displayed a wide range of size and shape in developing germ cells. A sub-population of elongated, tubular mitochondria in region 1 undergo a drastic morphological change, transitioning to dispersed, rounded organelles in region 2A through Fis1 and Drp1 mediated fission. The mitochondrial fragmentation, together with the lack of mtDNA replication in region 2A, effectively segregates mitochondrial genomes. The chance of potential complementation among different mitochondrial genomes is minimized. At region 2A, when mtDNA segregation takes place, mtDNA is not actively transcribed. Progressing into region 2B, mtDNA expression is activated, which triggers the biogenesis of ETCs and the activation of mitochondrial respiration. In heteroplasmic germ cells, mtDNA expression acts as a stress test for the integrity of mitochondrial genome. Mitochondria harboring wild-type genome will have functional ETCs and active respiration, whereas mitochondria containing deleterious mutations have defective ETCs and impaired respiration. At region 2B, mtDNA replication resumes, and preferentially takes place in healthy mitochondria (Hill et al., 2014). As a result, the proportion of wild-type genome increase through oogenesis. It is known that many nuclear-encoded mitochondrial proteins including several key factors required for mtDNA replication, are synthesized locally on mitochondrial surface by cytosolic ribosomes through a mitochondrial outer membrane protein Mdi (Zhang et al., 2016). This local translation allows coupling of synthesis and import of mitochondrial proteins. The import of preproteins across mitochondrial inner membrane requires mitochondrial membrane potential, which depends on active mitochondrial respiration (Geissler et al., 2000). We also found that local translation also preferentially takes place on healthy, polarized mitochondria (Zhang et al., 2019). Therefore, unhealthy mitochondria, due to the impaired local translation and import, will be starved of nuclear-encoded factors that are essential for mitochondrial biogenesis and mtDNA replication. This may underlie, or at least contribute to, the selective replication of wild-type genomes.

**Figure 6.**
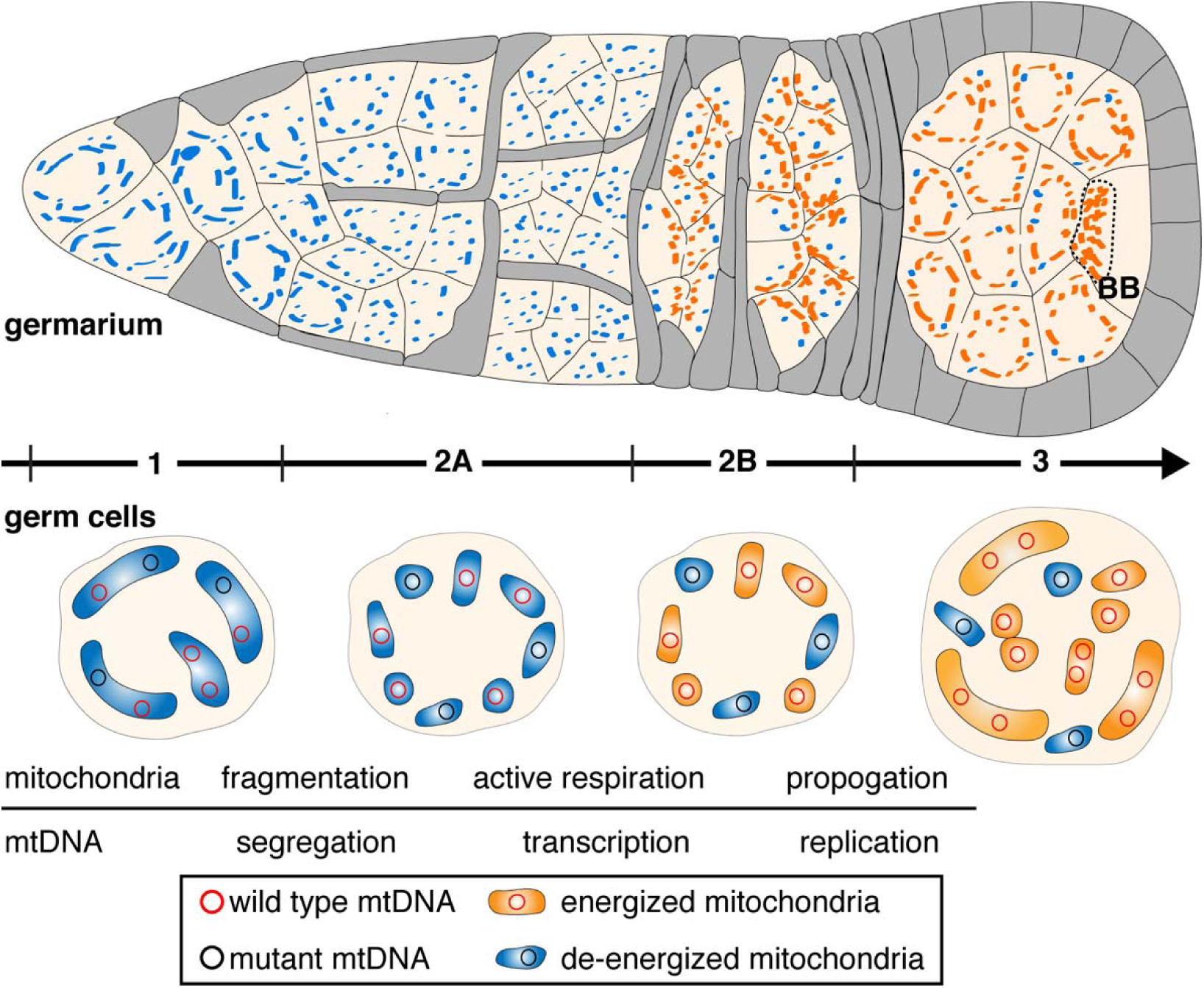
Developmentally-orchestrated mitochondria and mtDNA processes in the germarium of *Dm* are essential to limit the transmission of deleterious mutations. We propose that the mtDNA purifying selection is a developmentally regulated process. Mitochondrial fragmentation promotes mtDNA segregation in early germarium and prepare for effective selection on the organelle level based on the functional readout of the mtDNA within. Mitochondrial respiration is boosted in 16-cell stage region 2B, revealing the phenotype of the mtDNA and allowing selection based on mitochondrial functionality. In late germarium stage, healthy mitochondria containing wild-type mtDNA propagate much more vigorously than organelles containing deleterious mutations. These coordinated events act synergistically to secure the transmission of functional mtDNA from the female germline to the embryo. The Balbiani body (BB) in the oocyte further contributes to selective inheritance by concentrating wild-type mitochondria.

We previously found that mtDNA replication commences in the 16-cell cysts at region 2B, and is dependent on mitochondrial respiration. Hence, we proposed a model of selective inheritance through replication competition, in which healthy mitochondria containing a wild-type genome proliferate much more vigorously and outcompete these harboring deleterious mutations. This model explains the gradual decline of the load of deleterious mutations over generations. In mouse models, mutations on protein-coding genes that severely affect the mitochondrial respiratory chain activity were eliminated much faster than mild mutations on tRNA genes (Stewart et al., 2008a; Stewart et al., 2008b). This observation is consistent with our model of selection based on the functionality of individual genome, which might represent a conserved mechanism guiding mitochondrial inheritance in metazoans.

We found that a proportion of healthy mitochondria in the oocytes is pre-selected to join the Balbiani body, which eventually populates germ plasm in developing embryos. Thus, the Balbiani body enforces another purifying selection to further enhance mitochondrial fitness specifically in germ cells of offspring. The exact mechanism of how the healthy mitochondria are predetermined and unevenly distributed in the oocyte is unclear. Given the essential role of Milton in the formation of Balbiani body and mtDNA selective inheritance, it is possible that the healthy mitochondria are preferentially transported along the microtubules to Balbiani body, and then localized to the posterior end of oocytes. Balbiani body is a conserved structure in developing oocytes of several different species (Cox and Spradling, 2003; Pepling et al., 2007; Tworzydlo et al., 2016; Zhou et al., 2010). Its role in mitochondrial inheritance, particularly selective inheritance against damaging mtDNA mutations, remains to be explored in flies and other organisms.

Other models have also been proposed to explain how harmful mtDNA mutations are restricted from transmission through the female germline. Depolarized mitochondria can be cleared through Parkin-PINK1 mediated mitophagy in cultured cells (Narendra et al., 2010). A recent study demonstrated that mitophagy proteins ATG1 and BINP3 were required for mtDNA selection (Lieber et al., 2019). However, neither Parkin (Ma et al., 2014), nor ATG-8 dependent mitophagy involves in selective inheritance (Lieber et al., 2019; Zhang et al., 2019). This new type of selective mitophagy may act in parallel with the replication competition, to limit the transmission of deleterious mtDNA mutations. However, this potential redundancy cannot explain the complete loss of selection in the *mdi* mutant that lacks mtDNA replication in ovaries specifically (Zhang et al., 2019), but is largely healthy otherwise (Zhang et al., 2016). It is possible that Mdi, an outer-membrane has an unnoted role in the BINP3-mediated mitophagy, which awaits for future investigation.

Random segregation of mtDNA, enforced by a mitochondrial bottleneck, in principle could lead to the elimination of unhealthy germ cells or individuals with an excess of damaging mtDNA mutations through Darwinian selection (Stewart and Larsson, 2014). In *Drosophila*, mtDNA copy number remains constant from PGCs in embryo to GSCs and cystoblasts in pupae (Hurd et al., 2016). In the adult germaria region 2A, the absence of mtDNA replication (Hill et al., 2014), greatly reduces mtDNA copy number per germ cell at the 16-cell stage. However, all mitochondria from the 16-cell cyst derived from a single cystoblast end up in the oocyte (Ganguly et al., 2012). Thus, the lack of mtDNA replication in proliferating cysts should not be mistakenly considered as a way to generate bottleneck. We estimate each GSC contains ∼80 nucleoids. This number is in the range of the calculated value of mitochondria segregation unit (Ma et al., 2014). The relatively small number of mtDNA segregation unit, together with the large population size would be sufficient to facilitate the effective segregation of mitochondrial variants in *Dm*. For mtDNA mutations that do not severely impair mitochondrial respiration, but only mildly affect other mitochondrial activities, such as reactive oxygen species metabolism or heat production (Wallace, 2005), may not be sensitive to the selective inheritance described in this study. Nonetheless, the random segregation of different mitochondrial haplotypes, and through their interactions with both nuclear genome and environmental factors would produce progeny with different fitness outcome. This would exert a natural selection on organismal level and hence figure important in mtDNA selection and evolution (Lajbner et al., 2018; Wallace, 2005).

## Methods

### *Drosophila* genetics

Flies were maintained on cornmeal medium at 25 °C, unless otherwise stated. *w*^*1118*^ was used as the wild-type control. Heteroplasmic *mt:CoI*^*T300I*^ flies were generated as described previously (Hill et al., 2014), and maintained at 18 °C, unless otherwise stated. Generation of the TFAM-GFP reporter lines were described previously (Zhang et al., 2016). The UASp-AOX transgenic fly was generated by subcloning the Ciona intestinalis alternative oxidase coding sequence into the pUASp vector (Rorth, 1998), followed by standard germline transformation procedures (Chen et al., 2015). UASp-mtGFP transgenic fly was generated by subcloning EGFP cDNA into the pUASp vector (Rorth, 1998), followed by standard germline transformation procedures. *Fis1* RNAi (BL#63027), *cox5A* RNAi (BL#58282), *mtSSB* RNAi lines (BL#50600), *milton* RNAi (BL#43173), X-linked mChFP-Rho1(BL#52280), *bam-gal4* (BL#80579), *nanos-gal4* (BL#4937) came from the Bloomington *Drosophila* Stock Center (Bloomington, IN). *Drp1* RNAi (v44156), *tamas* RNAi (v3135) were from Vienna *Drosophila* Resource Center (VDRC). Vasa-GFP (#109171) came from the Kyoto *Drosophila* Genomics and Genetics Resources.

*Dm* fecundity and embryo hatch rate were performed according to previous study (Zhang et al., 2016). Knocking down of *Fis1, Drp1* driven by *bam-gal4*, knocking down of *cox5A, mtSSB, tamas* driven by *nanos-gal4*, and the overexpression of AOX by *nanos-gal4* did not change the fecundity of the female flies, as well as the hatch rate of their progeny (Fig. S5).

### Immunostaining of *Drosophila* germ cells

For staining of adult ovaries, 1-to-2-day old females were fed with yeast overnight prior to analysis. Ovaries were dissected in Schneider’s medium supplemented with 10% fetal bovine serum (FBS, Gibco) at room temperature. Ovaries were fixed for 20 minutes in 3.7% paraformaldehyde (Electron Microscopy Sciences) in PBS, then permeabilized in PBS containing 0.5% Triton-X100. After blocking in the PBSBT buffer (1 x PBS, 0.1% Triton-X100, 0.2% bovine serum albumin, BSA), the ovaries were incubated with primary antibodies overnight at 4 °C. Following washing with PBSBT buffer for three times, the ovaries were incubated with secondary antibodies at room temperature for 1 hr and washed again with PBSBT buffer for three times. Each ovariole was separated with fine-nose forceps under the stereomicroscope and mounted with Prolong Glass antifade mounting medium (Invitrogen) on the slides. Regular confocal imaging were collected on a Perkin Elmer Ultraview system and processed with Volocity software.

Antibodies used were as follows: mouse ATP synthase subunit α (Abcam, 15H4C4, 1:1,000), Rat α-Vasa (Developmental Studies Hybridoma Bank, DSHB, 1:200), Hts-1B1 (DSHB, 1:200); Alexa Fluor 647-Phalloidin (Invitrogen, 1:50); Alexa Fluor 568 goat α-Rat IgG (Invitrogen, 1:200), Alexa Fluor 568 goat α-mouse IgG (Invitrogen, 1:200), Alexa Fluor 568 goat α-rabbit IgG (Invitrogen, 1:200).

### Stimulated Emission Depletion (STED) microscopy and imaging quantification

STED microscopy equipped with a Leica 100x (1.4 N.A.) STED White objective was used for imaging germaria stained for ATP synthase subunit α and TFAM-GFP (Combs et al., 2019). 3D stack images (10 μm of z-stack and 0.16 μm/z-step) were analyzed with ImageJ. Cells from different germarium regions were selected manually by “freehand selection” function and isolated by “duplicate” function. Fluorescence outside the selected cell regions was removed by “clear outside” function. Background fluorescence value from each cell image was determined from an 1.5 × 1.5 μm^2^ region in the nucleus and subtracted from the original cell image by using “math-subtract” function. Individual mitochondrion and nucleoid from ATP syn α and TFAM-GFP channels, respectively, were called out using “color threshold” function with a “MaxEntropy” thresholding setting. Fluorescence outside the outlined ATP syn α and TFAM-GFP regions was removed by “clear outside” function. Next, we utilized an “restore selection” function to copy the outlined TFAM-GFP particle regions into the corresponding ATP syn α image. Then, the nucleoid number per mitochondrion was calculated manually as the number of outlined TFAM-GFP particles in an outlined ATP syn α particle. TFAM-GFP that marks nucleoids does not always displayed round dot shape. Instead, some puncta showed connection in between to form peanut shape staining, which may reflect different compaction state of mtDNA. To address this, 8-bit binary images for TFAM-GFP staining were generated using the “Make binary” function and further refined using a “watershed” function integrated in ImageJ to separate “touching” TFAM-GFP objects. After background removal and “clear outside” processing, non-mitochondria areas has zero fluorescence intensity, thus single mitochondrion was defined as ATP synthase subunit α staining with continuous pixels of fluorescence intensity. We noticed that the ATP synthase α subunit staining was not always uniform, with less intensity in the regions where nucleoids were located, which could be due to lack of cristae structures surrounding nucleoids (Stephan et al., 2019). Thus, a single mitochondrion was called if two adjacent, but distinct ATP synthase α puncta were connected with a single nucleoid. We found that there were ∼10% TFAM-GFP puncta in the cell localized in clustered mitochondria, which are unable to be individualized by ImageJ programs. This population of TFAM-GFP puncta were excluded from quantification.

### 3D Volume FIB-SEM

FIB-SEM were performed as previously described with modifications (Bleck et al., 2018). Ovaries were dissected in Schneider’s medium supplemented with 10% fetal bovine serum (FBS, Gibco), and immediately fixed in fixation solution (2.5% glutaraldehyde, 2% formaldehyde, 2 mM calcium chloride in 0.1 M sodium cacodylate buffer) at room temperature for 5 min, followed by an additional fixation on ice for 3 hr. After washing in cold cacodylate buffer containing 2 mM calcium chloride, the ovaries were post-fixed with reduced 2% Os_2_O_4_ (reduced by 1.5% potassium ferrocyanide right before use) for 1 hr on ice. Following washing with water, the tissues were placed in the thiocarbohydrazide (TCH) solution for 20 min at room temperature. Then, the ovaries were fixed in 2% Os_2_O_4_ for 30 min at room temperature, *en bloc* stained with 1% uranyl acetate overnight at 4°C, and further stained with Walton’s lead aspartate solution for 30 min at 60°C. After dehydration with ethanol series, the samples were embedded in Durcupan ACM resin (Electron Microscopy Sciences, Hatfield PA).

Embedded samples were then faced with a trim tool 90 diamond knife (DiATOME, Switzerland) on a Leica UCF-7 ultramicrotome (Vienna, Austria) and sputter coated with palladium/gold with a thickness of 50 nm in an EMS 575X sputter coater (Electron Microscopy Sciences, Hatfield, PA). The samples were imaged using a ZEISS Crossbeam 540 FIB-SEM microscope (Carl Zeiss Microscopy GmbH, Jena, Germany). Platinum and Carbon pads were deposited over the region of interest, and the run was set up and controlled by Atlas 5 software (Fibics Incorporated). SEM images were acquired using an accelerating 1.5 keV with a 1.5 nA of beam current and the in-lens detector captured backscattered electrons. The milling was performed with a FIB operating at 30 keV with a beam current 700 pA. The slice thickness and imaging pixel size were set to 10 × 10 × 10 nm voxels. The total volume acquired per tissue sample was as follows: wild-type: 29.59 × 26.19 × 30.30 µm (XYZ); *Fis1* knockdown: 29.06 × 31.67 × 19.29 µm (XYZ). The milling was performed from the anterior tip of the germaria towards posterior.

### Segmentation of mitochondria and cell clusters

Image segmentation was completed on a desktop PC (Thinkmate, Waltham MA) running Windows 10 with Intel Xeon Gold 6254 3.10 GHz processors, 2.0 TB RAM and an NVIDIA Titan V 12 GB VRAM video card. We used Dragonfly software (Ver. 4.1; Object Research Systems, Montreal QC) to segment the mitochondria and cells, and to collect mitochondrial morphometric information. Segmentation of mitochondria was completed using the U-Net convolutional neural network within the deep-learning tool in Dragonfly (Ronneberger et al., 2015). We used six raw FIB-SEM images from evenly distributed regions (approx. one image every 400 steps) of the wild-type sample to build our mitochondrial training set for the U-Net model. Mitochondria were initially segmented by labelling the outer-mitochondrial membrane (OMM) via thresholding followed by manual clean-up. Next, the mitochondrial matrix was segmented using the fill inner area tool of the OMM region of interest (ROI) followed by the erode tool and manual cleaning to eliminate the OMM ROI. We used this training dataset to build the U-net model where training data were augmented (horizontal and vertical flip, 180° max rotation, 10° max shear, 75-150 % scale, 0-2 brightness, and 0-0.10 Gaussian noise) and 20% of the training data were used for validation. We used categorical cross-entropy for our loss function and Adadelta for our optimization algorithm. The model was then applied to the full volume to segment out the mitochondrial matrix, which was then manually cleaned.

To segment individual mitochondria, we used a watershed mapping approach. Watershed map seed points were isolated from the previously described mitochondrial matrix ROI using a connected components analysis. The boundaries of this watershed map were set by the OMM, which was included by dilating the mitochondrial matrix ROI – followed by manual cleaning. A minor population of segmented objects have light staining, which could be swollen mitochondria, undifferentiated mitochondria that have less cristae, or *Wolbachia*. We were unable to unequivocally call out *Wolbachia* based on the presence of three layers of membranes, a distinct feature of *Wolbachia* (White et al., 2017), due to the limited resolution of FIB-SEM. Nonetheless, we ran analyses on dense-stained objects only, and found the trend of mitochondrial fragmentation was essentially the same as shown in Figure 1, demonstrating that potential inclusion of *Wolbachia* does not interfere with our analyses. Importantly, Fis1 knockdown, which should have no impact on the morphology of *Wolbachia*, effectively shifted the morphology of objects, further supporting our conclusion.

In FIB-SEM images, the size of nuclei and locations of cells were used to distinguish germ cells from somatic cells. For germ cells, their relative locations in a germarium, and the number of interconnected germ cells, judged by presence of ring canals and connecting fusome within a cyst, were used to determine their developmental stage. Cell clusters were segmented from the 3D volume using z-interpolation over 50 image intervals. Cell cluster-specific mitochondrial surface area and volume were extracted from these datasets.

### Membrane potential staining

Adult ovaries were dissected in Schneider’s medium supplemented with 10% fetal bovine serum and incubated with medium containing Tetramethylrhodamine (TMRM, Invitrogen, 1:1000) and mitoTracker Green (Invitrogen, 100 nM) for 20 min. The ovaries were rinsed with PBS for 3 times, and then imaged live on a Perkin Elmer Ultraview system within 1 hr. The position and morphological characters were used to define developmental stages of germ cells. The 16-cell cyst in region 2B is flatten and extends the entire width of the germarium. Mitochondria are closely associated with fusome at this region, thus the mitochondrial staining within the cyst at this stage display a long, branched appearance along the fusome structures. Region 2A cysts are considered as locating immediately anterior of the first region 2B cyst.

### Mitochondrial activity staining and EdU labelling of *Drosophila* adult ovaries

Histochemical staining for the activity of mitochondrial succinate dehydrogenase (complex II) and cytochrome C oxidase (complex IV) in ovaries was performed as previously described (Wang et al., 2019). EdU incorporation assay in *Drosophila* ovaries was carried out as described (Hill et al., 2014). The ovaries were co-stained for fusome to define developmental stages of germ cells.

### Single-molecule fluorescence *in situ* hybridization and quantification

The mitochondrial and nuclear-encoded transcripts in the germarium was detected using a single-molecule fluorescence *in situ* hybridization (smFISH) protocol published previously (Trcek et al., 2017). The sequence of fluorescently labelled short DNA probes targeting to *ND4, cox1, NDUFB5, cox5A, Fis1, Drp1, mtSSB* mRNA are listed in Table S2.

Series images with FISH and nuclear (DAPI) channels were collected by an Instant Structured Illumination (iSIM) microscope (VisiTech International) and processed in ImageJ. Germline cells and germarium regions were identified based upon the position, morphological characters and the nuclei size (DAPI staining). Nuclei of region 2B germ cells are round and surrounded by pre-follicle cells that have smaller oval or round shape nuclei. Region 2A cysts are located immediately anterior of the first region 2B cyst. To quantify relative mRNA level of *ND4, NDUFB5, COX1*, or *COX5A* in the region 2A and 2B cysts (Figure 3B), cysts from germarium region 2A and 2B were manually outlined, duplicated to display as two different images with the “Duplicate” function, and isolated by the “clear outside” function, respectively. The “3D objects counter” plugin with smallest threshold setting was used to select whole cyst region and to quantify cyst volume from each series image. The “3D objects counter” plugin with automatic threshold setting was used to select and to quantify fluorescence density of each FISH punctum in 3D. Background fluorescence value was subtracted from total FISH fluorescence density in each germarium region. The resulting net FISH fluorescence density was divided by the corresponding cyst volume.

RNAi for *Fis1, Drp1, cox5A*, and *mtSSB* were performed specifically in the germ cell. The abundance of these transcripts in the follicle cells should not be affected. Therefore, we utilized the smFISH fluorescence intensity in follicle cells as an internal control to determine the knockdown efficiency of each RNAi line. In ImageJ, the areas of region 2A cysts (for *Fis1* and *Drp1* RNAi) or region 2B cysts (for *cox5A* and *mtSSB* RNAi), and their corresponding region 2B follicle cells, were manually outlined. The fluorescence intensity from each selected region was calculated with the method describe above. The resulting smFISH fluorescence intensity from region 2A or region 2B cysts were normalized by that of the corresponding region 2B follicle cells.

### Primordial germ cell isolation from *Dm*

The primordial germ cells from *Dm* embryos and pupae were isolated as previously described (Shigenobu et al., 2006) with modification. A Vasa-GFP transgene that specifically labels germ cells throughout the life cycle was used to isolate the germ cells using fluorescence-activated cell sorting (FACS) assay. Female PGCs were separated from male PGCs by an X-chromosome linked monomeric cherry fluorescence (mChFP) tagged Rho1 protein under control of Rho1 regulatory sequence (Abreu-Blanco et al., 2014). The female Vasa-GFP transgenic flies, which carry wild-type or heteroplasmic *mt:CoI*^*T300I*^ mtDNA, were crossed with male X-linked mChFP-Rho1 flies in cages and allowed to lay eggs on a grape agar plate (Genesee Scientific. Inc). The germ cells of the female progeny will carry both GFP and mChFP fluorescence. Following pre-collection for 3 hr, the embryos were collected and allowed to develop till stage 15 at 25 °C (staging according to (Williamson and Lehmann, 1996)). The embryos were then dechorionated for 30 s in 50% bleach. After washing with water, the embryos were transferred to a microcentrifuge tube filled with 500 µl of Schneider’s insect medium (Gibco). The blue pestle matching with the microcentrifuge tube (USA Scientific, Inc.) was used to gently homogenize the embryos. The homogenate was filtered through a 125 µm mesh and then centrifuged at 860 g for 1 min at 4 °C. After one wash in ice-cold calcium-free Schneider’s medium (Sigma), the pellet was resuspended in calcium-free Schneider’s medium containing 0.25% trypsin (Invitrogen) and incubated at 37°C for 10 min. The cell suspension was filtered through a 35 μm mesh, and the same amount of Schneider’s medium supplemented with 20% fetal bovine serum was added to stop the trypsinization. The dissociated cells were pelleted by centrifugation at 860 g for 1 min. The cells were resuspended in Schneider’s medium and filtered through a 35 μm mesh immediately before cell sorting. Flow cytometry analyses were performed on a BD FACSCalibur flow cytometer and analyzed with FACSDiva. The female PGCs were sorted by gating for GFP and mChFP-positive events. Female somatic cells were sorted by gating for GFP-negative and mChFP-positive events. The quality and purity of the sorted PGCs were confirmed by fluorescence microscope. For isolation of PGCs from pupae, the staged embryos were transferred to the standard cornmeal medium till the desired development stage. All the other procedures were the same except for the exclusion of the dechorionation step.

### *Drosophila* ovaries stem cell culture

The fGS/OSS is a stable cell line consisting of a mixture of *Dm* adult female germline stem cell (fGS) and ovarian somatic sheath cells (OSS). OSS is the derivative line from which the fGS component has been lost. Both cell lines were obtained from *Drosophila* Genomics Resource Center (DGRC) and cultured as described previously (Niki et al., 2006).

### Measurement of mtDNA copy number

To quantify mtDNA copy number, total DNA was isolated from the FACS-sorted PGCs or the somatic cells using QIAamp DNA Micro Kit (Qiagen). The mtDNA copy number was measured using droplet digital PCR (ddPCR, Bio-rad), a method for absolute quantification of nucleic acids. Primers were targeted to the mtDNA-encoded cytochrome c oxidase subunit I (CoI), and the nuclear-encoded Histone 4 (His4) genes. Primers and probes used in ddPCR are as follows: CoI-forward: 5’ATTGGAGTTAATTTAACATTTTTTCCTCA3’, CoI-rev: 5’AGTTGATACAATATTTCATGT-TGTGTAAG3’, CoI-probe: 5’AATACCTCGACGTTATTCAGATTACCCA3’, His4-for: 5’TCCAAGGTATCA-CGAAGCC3’, His4-rev: 5’AACCTTCAGAACGCCAC3’, His4-probe: 5’AGCGCATATCTGGACTCATATACGAG3’. The CoI-probe and His4-probe were synthesized by labeling the 5’ nucleotide with FAM and HEX reporter fluorophores, respectively. Around 1 µg of total DNA was digested with EcoRI at 37 °C for 1 hr. Then the ddPCR reaction mixture was assembled to contain 1x ddPCR Supermix for probes (Bio-rad), 900 nM of each forward and reverse primers, 250 nM of probe, and up to 1 ng of total DNA. The reaction was conducted in the QX200™ Droplet Generator, followed by the thermal cycler and analyzed by the QX200 Droplet reader as per the manufacturer’s instruction.

### mtDNA selection and quantification of heteroplasmy

mtDNA selection in the female germline was carried out according to previous study (Zhang et al., 2019). Briefly, heteroplasmic flies with wild-type and knockdown nuclear backgrounds were transferred from 18°C to 29°C right after eclosion. Individual heteroplasmic female fly was mated with five male *w*^*1118*^ flies at 29°C and at least ten female flies were analyzed. Eggs produced during the first 6 days were discarded. The eggs laid on the 7^th^ day were pooled and the heteroplasmic levels were compared between mother and her eggs. Quantification of heteroplasmic mtDNA was performed as described previously (Hill et al., 2014).

### Statistical analysis

Data were analyzed using Student’s *t* test, Mann-Whitney test or one-way analysis of variance. The difference was considered statistically significant when *p* < 0.05.

## Acknowledgements

We thank F. Chanut for comments and edits on the manuscript; B. Glancy for advice and comments on the FIB-SEM work; M. Aronova and J. Cohen for EM technical assistance; NHLBI FACS core and NCI CCR Genomics core for technical support; Bloomington *Drosophila* Stock Center, Vienna *Drosophila* Resource Center, Kyoto *Drosophila* Genomics and Genetics Resources for fly stocks; *Drosophila* Genomics Resource Center for *Dm* stem cell cultures; Developmental Studies Hybridoma Bank for antibodies. This work is supported by NHLBI Intramural Research Program.

## Supplemental Materials

### Supplementary Tables

**Table S1.** Frequency distribution of mitochondrial volume in control and *Fis1* knockdown (driven by *bam-gal4*) germarium region 1 and region 2A. The data are percentage (%).

| Volume ($\mu\text{m}^3$) | ctrl | | *Fis1*-KD | |
| --- | --- | --- | --- | --- |
|  | region 1 | region 2A | region 1 | region 2A |
| <0.01 | 10.3 | 13.5 | 12.1 | 16.1 |
| 0.01-0.03 | 27.2 | 50.0 | 19.7 | 35.1 |
| 0.03-0.05 | 14.7 | 17.7 | 15.6 | 18.2 |
| 0.05-0.07 | 9.4 | 4.5 | 11.4 | 7.4 |
| 0.07-0.09 | 7.4 | 1.9 | 7.6 | 3.9 |
| 0.09-0.11 | 6.2 | 2.4 | 5.1 | 3.2 |
| 0.11-0.13 | 5.7 | 2.4 | 8.6 | 4.2 |
| 0.13-0.15 | 5.2 | 2.0 | 6.9 | 3.5 |
| 0.15-0.17 | 6.3 | 2.2 | 4.1 | 2.3 |
| 0.17-0.19 | 3.8 | 1.8 | 2.7 | 2.0 |
| 0.19-0.21 | 2.1 | 1.1 | 1.7 | 1.6 |
| 0.21-0.23 | 1.2 | 0.4 | 2.5 | 1.1 |
| 0.23-0.25 | 0.4 | 0.1 | 1.2 | 0.7 |
| 0.25-0.27 | 0.1 | 0 | 0.4 | 0.4 |
| >0.27 | 0 | 0 | 0.4 | 0.3 |

**Table S2.** The sequence of fluorescence *in situ* hybridization short DNA probes targeting to *ND4, cox1, NDUFB5, cox5A, Fis1, Drp1, mtSSB* mRNA.

**Video 1.** 3D FIB-SEM image stack of wild-type *Dm* early germarium with automated mitochondrial segmentation. Y-stack images are shown and time represents sequential images moving across the depth of the germarium (10 nm steps). Germline cysts are labelled with different colors.

**Video 2.** 3D FIB-SEM image stack of *Fis1* knockdown *Dm* germarium with automated mitochondrial segmentation.

**Supplementary Figure 1.**
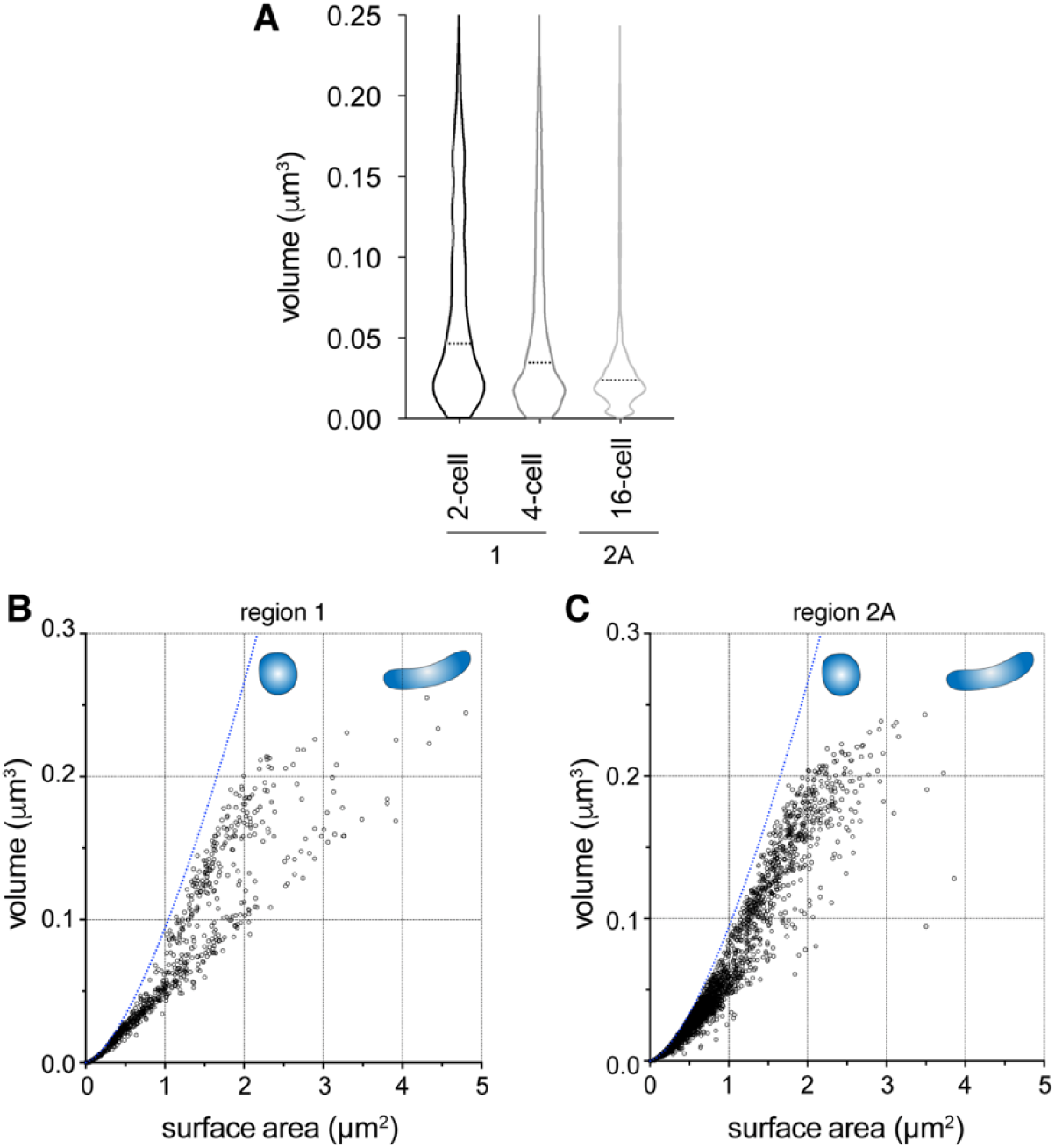
FIB-SEM analyses show the change of mitochondrial volume and morphology in early germarium. **(A)** A violin plot showing the distribution of individual mitochondrial volume in germarium 2-cell, 4-cell and 16 cell cysts. The dashed lines indicate the median volume. **(B and C)**, The volume (V) versus surface area (SA) of each mitochondrion from region 1 **(B)** and region 2A **(C)** in wild-type ovaries. The dashed blue lines represent the relationship between V and SA of perfect spheres, which have the lowest SA/V ratio among all shapes. For a given volume, the lower SA/V suggests more rounded shape. Please be noted that more mitochondria in region 2A shift towards the blue line, compared with those in 2-cell cyst at region 1.

**Supplementary Figure 2.**
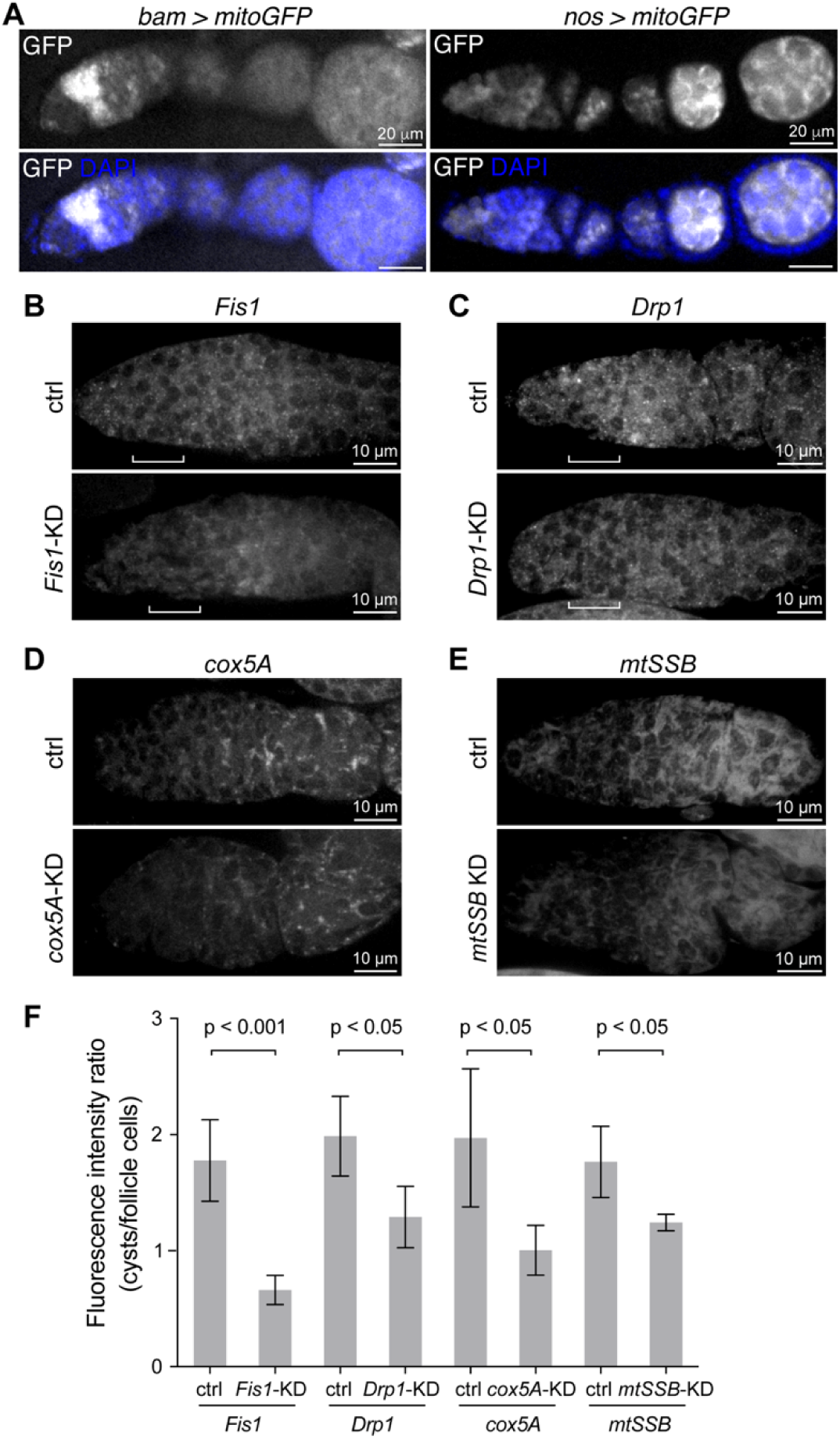
Expression pattern of germline specific drivers and FISH assay. **(A)** Expression pattern of mitochondrially-targeted GFP (mtGFP) driven by *bam-gal4* and *nanos-gal4* drivers. Note the high expression level of mtGFP in dividing cysts driven by *bam-gal4*. Ovaries are co-stained with DAPI (blue). Scale bar, 20 µm. **(B-E)** smFISH assay show the knockdown efficiency of several RNAi lines used in this work. The images are smFISH using DNA probes targeted to *Fis1* **(B)**, *Drp1* **(C)**, *cox5A* **(D)**, *mtSSB* **(E)** RNA in ovaries. Brackets in **(B)** and **(C)** indicate region 2A. Scale bar, 10 µm. (F) Quantification of the smFISH fluorescence intensity in germline cysts using region 2B follicle cells of the same sample as the internal control. The ratio of the fluorescence intensity in germline cysts to follicle cells are shown.

**Supplementary Figure 3.**
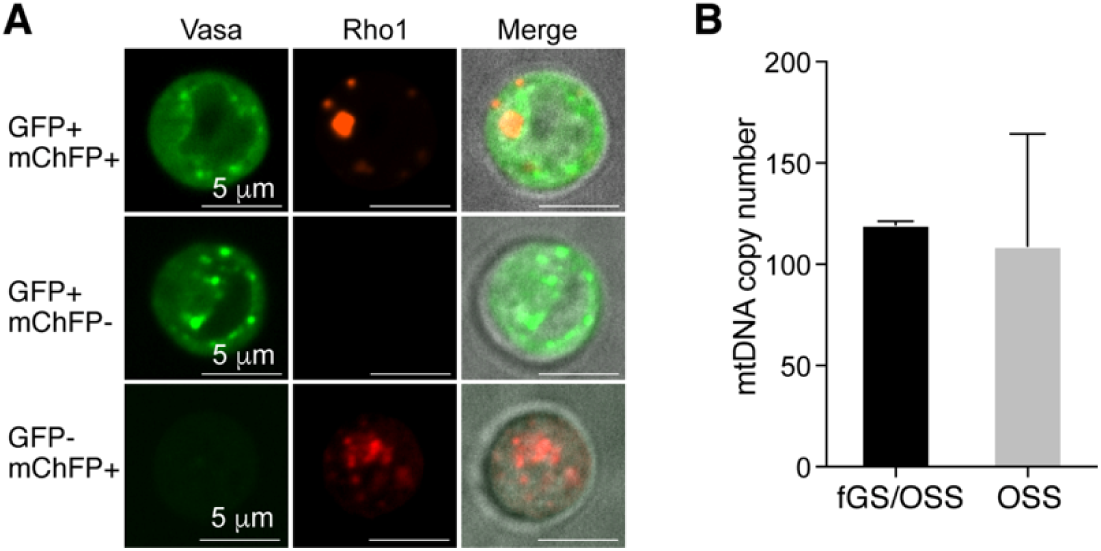
Quantification of mtDNA copy number in GSCs. **(A)** Confocal imaging of cells isolated from a mix of female Rho1-mChFP/X; Vasa-GFP/+ and male Y/X; Vasa-GFP/+ early pupae using FACS. The germ cells of the female pupae carry both GFP and mChFP fluorescence. The merged images are overlay of GFP and mChFP fluorescence with DIC channel. Scale bar, 5 µm. **(B)** mtDNA copy number per cell in fGS/OSS (the mixture of female germline stem cell and ovarian somatic sheath cells) and OSS (ovarian somatic sheath) cells quantified by real time PCR. The average copy number of mtDNA in fGS/OSS are 119.5 and 109, respectively. Thus, we deduce the mtDNA number in fGS alone should be similar to that of fGS/OSS.

**Supplementary Figure 4.**
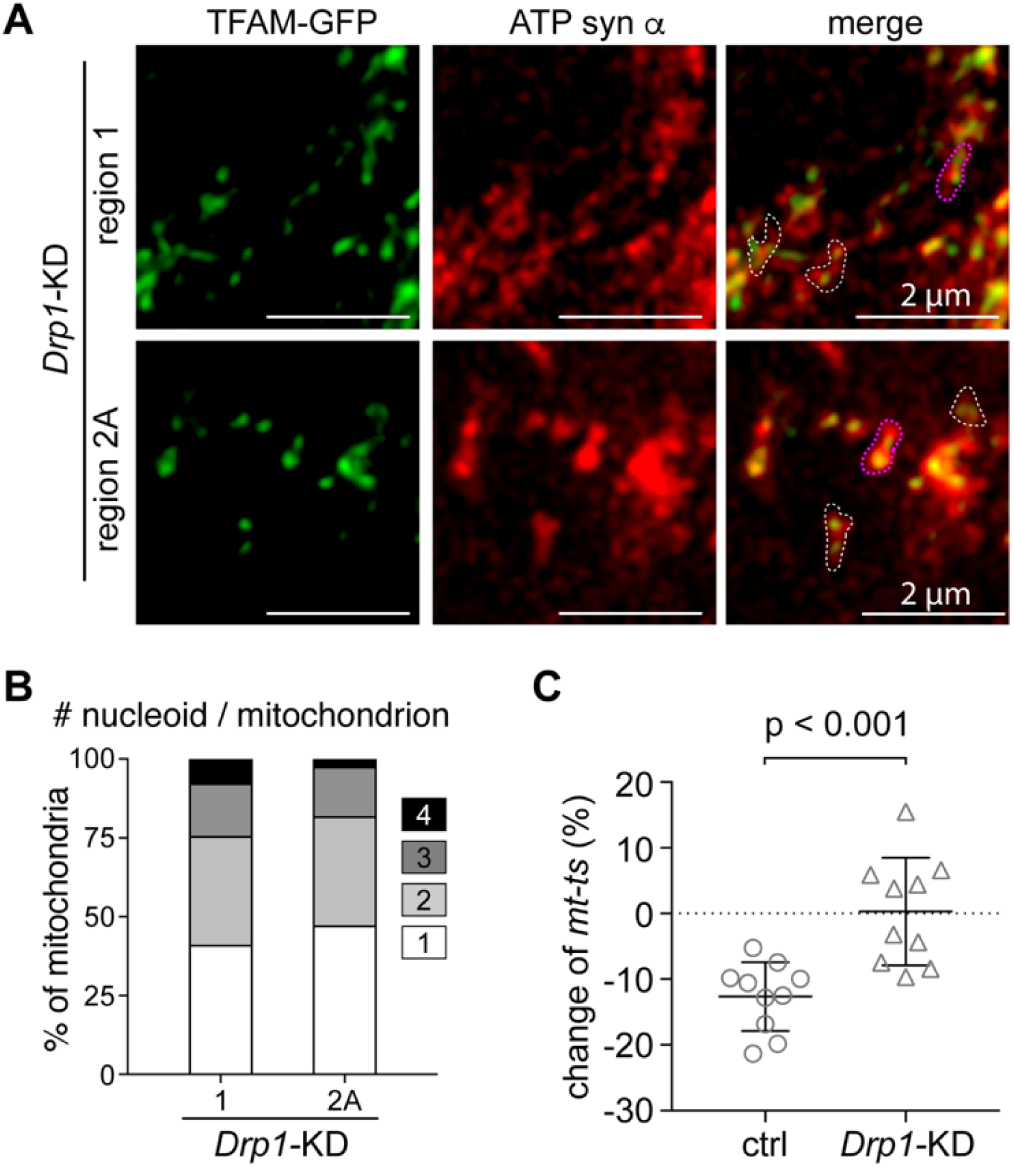
*Drp1* knockdown impairs nucleoid segregation and mtDNA selective inheritance. **(A)** Representative images of the anterior end of region 1 and region 2A in a *Drp1* knockdown germarium (driven by *bam-gal4*) labeled with TFAM-GFP and ATP synthase ⍰-subunit. An object with continuous ATP synthase ⍰-subunit staining was defined as a single mitochondrion (white dashed lines). Two adjacent, but distinct ATP synthase subunit α puncta were also called as a single mitochondrion if they appeared in the same contour and were connected with one nucleoid (magenta dashed lines). Scale bar, 2 µm. **(B)** The number of nucleoids per mitochondrion was determined by counting the number of TFAM-GFP puncta within a single mitochondrion shown in **(A)** in regions 1 (stem cells or cystoblasts) and 2A from *Drp1* knockdown ovaries. The fraction of each group is shown (n = 10 cysts for each group). **(C)** Knockdown of *Drp1* using *bam-gal4* driver diminishes the selection against the *ts* mtDNA in heteroplasmic *mt:CoI*^*T300I*^ *Drosophila* (*p*<0.001).

**Supplementary Figure 5.**
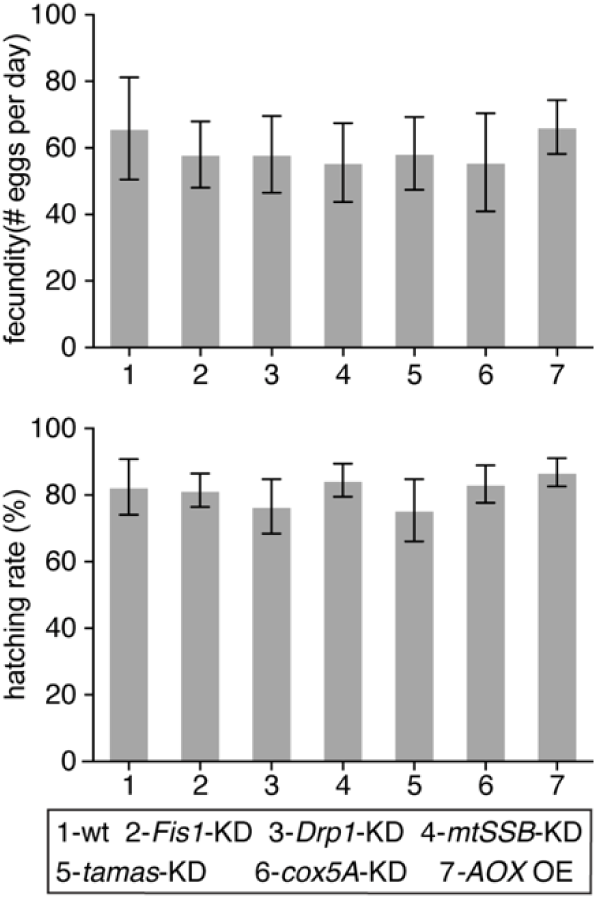
*Drosophila* fecundity and hatching rate. The fecundity and hatching rate of their progeny in *Fis1, Drp1* knockdown (driven by *bam-gal4*), *cox5A, mtSSB, tamas* knockdown (driven by *nanos-gal4*), and the AOX ectopic expression (driven by *nanos-gal4*) flies are comparable with those in wild-type flies.

**Supplementary Figure 6.**
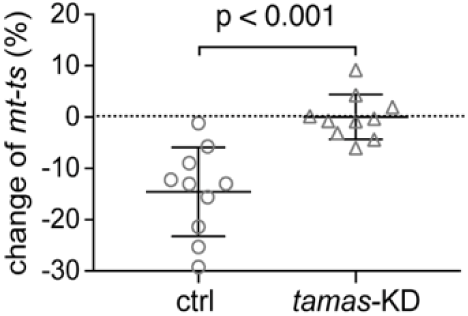
Knockdown of *tamas* in ovary diminished mtDNA selection. Knockdown of *tamas* using *nanos-gal4* driver diminishes the selection against the *ts* mtDNA in heteroplasmic *mt:CoI*^*T300I*^ *Drosophila* (*p*<0.001).

